# Quantitative analysis of cryptic splicing associated with TDP-43 depletion

**DOI:** 10.1101/076117

**Authors:** Jack Humphrey, Warren Emmett, Pietro Fratta, Adrian M. Isaacs, Vincent Plagnol

**Affiliations:** University College London Genetics Institute, Gower Street, London WC1E 6BT, UK; Department of Neurodegenerative Disease, UCL Institute of Neurology, Queen Square, London WC1N 3BG, UK; Department of Molecular Neuroscience, UCL Institute of Neurology, Queen Square, London, WC1N 3BG, UK; Sobell Department of Motor Neuroscience and Movement Disorders, UCL Institute of Neurology, Queen Square, London WC1N 3BG, UK; The Francis Crick Institute, 1 Midland Road, London NW1 1AT, UK

## Abstract

Reliable exon recognition is key to the splicing of pre-mRNAs into mature mRNAs. TDP-43 is an RNA-binding protein whose nuclear loss and cytoplasmic aggregation are a hallmark pathology in amyotrophic lateral sclerosis and frontotemporal dementia (ALS/FTD). TDP-43 depletion causes the aberrant inclusion of cryptic exons into a range of transcripts, but their extent, relevance to disease pathogenesis and whether they are caused by other RNA-binding proteins implicated in ALS/FTD are unknown. We developed an analysis pipeline to discover and quantify cryptic exon inclusion and applied it to publicly available human and murine RNA-sequencing data. We detected widespread cryptic splicing in TDP-43 depletion datasets but almost none in another ALS/FTD-linked protein FUS. Sequence motif and iCLIP analysis of cryptic exons demonstrated that they are bound by TDP-43. Unlike the cryptic exons seen in hnRNP C depletion, those linked to TDP-43 do not originate from a particular family of transposable element. Cryptic exons are poorly conserved and inclusion overwhelmingly leads to nonsense-mediated decay of the host transcript, with reduced transcript levels observed in differential expression analysis. RNA-protein interaction data on 73 different RNA-binding proteins showed that, in addition to TDP-43, 7 specifically bind TDP-43 linked cryptic exons.This suggests that TDP-43 competes with other splicing factors for binding to cryptic exons and can repress cryptic exon inclusion. Our quantitative analysis pipeline confirms the presence of cryptic exons during TDP-43 depletion and will aid investigation of their relevance to disease.

## 1 Introduction

Splicing depends on the reliable recognition and subsequent removal of non-coding intronic sequence by the multi-protein spliceosome complex. Boundaries that demarcate exons from introns are first recognised by the binding of the U1 small nuclear RNA (snRNA) to the 5´ splice site at the beginning of the intron and the U2 snRNA binding to the polypyrimidine tract and 3´ splice site at the end of the intron. The two snRNAs then interact with each other and the intronic sequence is removed via a series of transesterification reactions (Matera et al., 2014). During splicing, regulatory sequences in the nascent RNAs recruit RNA-binding proteins (RBPs) to shape mRNAs, selecting or omitting particular exons by acting to enhance or repress splicing.

Due to the long length and reduced evolutionary conservation of intronic sequences, pairs of 3′ and 5′ splice sites can emerge randomly to create potentially new exons. These cryptic exons (also known as pseudoexons) arise due to mutations that create new splice sites or remove the existing binding sites for splicing repressors. These type of mutations have been also implicated in in a number of genetic diseases (Buratti et al., 2007; Eng et al., 2004; Meili et al., 2009; Vorechovsky, 2006). Inclusion of a cryptic exon, untested by evolution, can destabilise the transcript or radically alter the eventual protein structure. The former can occur by the nonsense mediated decay (NMD) pathway, which occurs when a transcript has a premature termination codon introduced either directly by the cryptic exon or by a subsequent shift of reading frame (McGlincy and Smith, 2008).

Cryptic exons can also emerge from transposable elements. One such example are Alu elements, the predominant transposable element in primates which are often found within introns in the antisense direction (Deininger and Prescott, 2011). The consensus Alu sequence consists of two arms joined by an adenine-rich linker ending with a poly-adenine tail. When transcribed in the antisense direction these uridine-rich sequences can act as cryptic polypyrimidine tracts and only a few mutations are required to convert them into viable exons in a process termed exonisation (Sorek et al., 2002). De novo mutations that lead to Alu exonisation have been found in a range of diseases (Vorechovsky, 2010) suggesting a need for regulation of potentially damaging Alu exons. Alu exonisation is repressed by the RNA binding protein hnRNP C, which competes with the spliceosome component protein U2AF65, the partner of the U2 snRNA, for binding cryptic 3′ splice sites (Zarnack et al., 2013). Due to the potentially negative effects of incorporation of new exons, aberrant recognition of cryptic exons needs to be repressed.

TDP-43 is an RNA-binding protein encoded by the *TARDBP* gene. Loss of TDP-43 from the nucleus accompanied by TDP-43 positive inclusions in the cytoplasm of cortical and spinal cord neurons is the hallmark pathology of amyotrophic lateral sclerosis (ALS) as well as the majority of cases of frontotemporal dementia (FTD) (Neumann et al., 2006). In addition, missense mutations in *TARDBP* can cause familial ALS (Sreedharan et al., 2008). These findings point to a central role of TDP-43 in the aetiology of ALS.

Rare ALS-causing mutations have been found in the genes coding for other RNA-binding proteins such as FUS, hnRNP A1 and hnRNP A2B1 (Kim et al., 2013; Vance et al., 2009) further raising the possibility that the impairment of RNA processing is a central cause of ALS. TDP-43 was first shown to repress the inclusion of exon 9 in the *CFTR* gene by binding to long UG-rich sequences (Buratti and Baralle, 2001) and subsequently shown to act as both a splicing enhancer and repressor (Bose et al., 2008; Mercado et al., 2005; Shiga et al., 2012). Transcriptome-wide studies of the effects of TDP-43 depletion, overexpression or mutation have demonstrated widespread changes in gene expression and splicing (Arnold et al., 2013; Chiang et al., 2010; Polymenidou et al., 2011; Shan et al., 2010). Of particular interest are long intron containing genes which are dramatically downregulated in TDP-43 depletion conditions.

Recently, Ling and colleagues observed the inclusion of cryptic exons when TDP-43 was depleted in HeLa or mouse embryonic stem cells (Ling et al., 2015). These cryptic exons were shown to originate from poorly evolutionarily conserved sequence and shared no positions between the two species. These findings raise the possibility that impaired exon recognition contributes to TDP-43’s role in ALS aetiology.

Given the intriguing possibility that impaired recognition of exons plays a causal role in ALS, we aimed at replicating and expanding the findings of Ling and colleagues. Firstly, we undertook a quantitative genome-wide analysis of cryptic exon patterns, defining objective criteria that take advantage of biological replicates when available. Secondly, we applied this computational strategy to seven datasets (four human and three murine models) to systematically quantify cryptic RNA alterations associated with depletion of TDP-43 as well as FUS and hnRNP C, two proteins linked to ALS and cryptic exons respectively. Lastly, we used independent protein-RNA interaction datasets, conservation data, repeat element annotation and splice site scoring to investigate the potential mechanisms linking TDP-43 depletion with the cryptic exon phenomenon.

**Table 1.** All RNA-sequencing data used in this study. ES: embryonic stem cell. K562: human leukaemia cell line. siRNA: small interfering RNA. shRNA: short hairpin RNA. ASO: antisense oligonucleotide. PE: paired end sequencing. SE: single end sequencing. For single end sequencing, depth is measured in millions of mapped reads whereas paired end sequencing depth is measured in millions of mapped fragments.

|  | Species | Cell Type | Protein | Depletion method | Library type | Read type | Depth | Citation |
| --- | --- | --- | --- | --- | --- | --- | --- | --- |
| 1 | Human | HeLa | TDP-43 | siRNA | mRNA | 100bp PE | 97-116M | Ling, 2015 |
| 2 | Mouse | ES | TDP-43 | Knockout | mRNA | 100bp PE | 70-75M | Ling, 2015 |
| 3 | Human | K562 | TDP-43 | shRNA | Total RNA | 100bp PE | 55-62M | ENCODE |
| 4 | Human | K562 | TDP-43 | shRNA | mRNA | 100bp PE | 25-29M | ENCODE |
| 5 | Mouse | Adult brain | TDP-43 | ASO | mRNA | 75bp SE | 35-60M | Polymenidou, 2011 |
| 6 | Mouse | ES | TDP-43 | Knockout | mRNA | 40bp SE | 2-11M | Chiang, 2010 |
| 7 | Human | K562 | FUS | shRNA | mRNA | 100bp SE | 12-21M | ENCODE |
| 8 | Mouse | Adult brain | FUS | ASO | mRNA | 72bp SE | 20-60M | Lagier-Tourenne, 2012 |
| 9 | Human | HeLa | hnRNP C | siRNA | mRNA | 72bp SE | 26-28M | Zarnack, 2013 |

## 2 Results

### 2.1 Depletion of TDP-43 but not FUS results in cryptic exons

We analyzed publicly available TDP-43 depletion RNA-Seq datasets (three human, three murine, datasets 1-6 in Table 1), FUS depletion RNA-Seq datasets (1 human, 1 murine, datasets 7-8 in Table 1) and as a positive control a human hnRNP C depletion dataset for which cryptic exons have previously been reported (dataset 9 in Table 1). While these datasets differ in library preparation method, read depth and length, and protein depletion method (Table 1), the FUS datasets match the TDP-43 datasets in cell type. Using these datasets, we took a genome-wide approach to describe the pattern of unannotated splicing in human and mouse that we call *CryptEx*. The first step involves identifying novel splice junctions between annotated exons and unannotated intronic regions. These reads were clustered to create putative cryptic exons. Making use of biological replicate samples we compared the abundance of reads covering the cryptic region relative to the rest of the annotated exons of the surrounding gene. Differential usage of each cryptic region was tested between depletion and control samples. Results were omitted if they fell outside of a strict 5% false discovery rate.

In order to further refine our cryptic exon detection algorithm, we then measured the amount of splicing to and from each region and used this information to filter and classify cryptic splicing events into three categories of cryptic exon: (i) cassette-like, where novel 3′ and 5′ splice sites are recognised forming a completely new exon; (ii) 5′ extension, where a novel 3′ splice site is recognised and an existing exon is extended upstream of its annotated start and (iii) 3′ extension, where a novel 5′ splice site is recognised and an exon is extended downstream of its annotated end (Figure 1A). Note that our approach does not consider fully retained introns, but several other methods have been designed for this purpose (Bai et al., 2015; Li et al., 2015; Pimentel et al., 2015).

**Figure 1.**
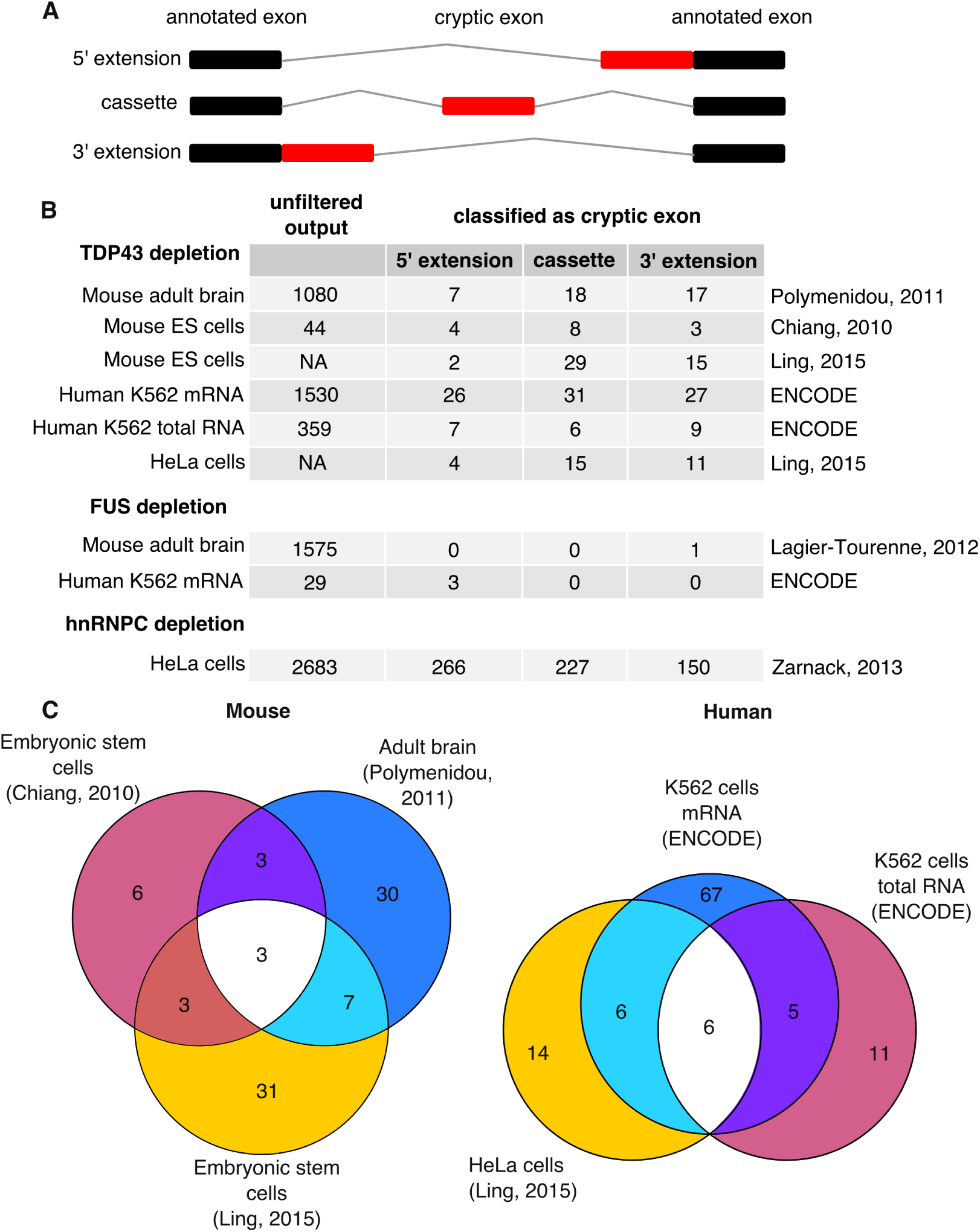
Cryptic splicing discovered by the *CryptEx* pipeline. A– Schematic of the three classes of cryptic exon. Black boxes represent annotated exons and red boxes represent a cryptic exon. Grey lines represent the spliced intron. B-Tally of the three classes of cryptic exon discovered by the Cryptex pipeline across the nine datasets. “Unfiltered output” refers to the number of differentially used cryptic splicing events at a false discovery rate (FDR) < 5% before undergoing cryptic exon classification. Counts from Ling et al’s data are taken from the paper itself. C-Venn diagrams showing the overlap between the six TDP-43 depletion datasets.

Figure 1B lists the counts of both pre-classification cryptic regions (“unfiltered output”) and the post-classification cryptic exons. Comparing the two human ENCODE K562 cell line TDP-43 depletion datasets (3-4), the poly-A selected mRNA-Seq dataset yielded far more splicing events than the total RNA dataset, presumably due to polyA selection leading to a higher coverage of mature spliced mRNA species. In total 95 human cryptic exons were discovered and classified, with the majority only detected in the mRNA-seq dataset. 11 cryptic splicing events were shared between datasets 3 and 4 (Figure 1C). Of the 26 human cryptic exons reported by Ling, 12 were seen in at least one of the two datasets 3 and 4.

Both mouse datasets differ in both cell type (adult striatum in dataset 5 vs embryonic stem (ES) cell in dataset 6) and read depth (35-60M in dataset 5 vs 2-10M in dataset 6). 52 cryptic exons were identified in total, with 46 detected in the adult striatum and 15 in ES cells, with 6 exons observed in both. Of the 46 cryptic splicing events identified in murine samples by Ling et al, 13 were detected in at least one of datasets 5 and 6. Side by side visual inspection (Figures S2 and S3) suggests that differences in library preparation and read depth are behind the low the concordance rates in both human and mouse, as cryptic exons detected in the higher depth dataset (K562 mRNA and mouse adult brain) can be observed by eye in the lower depth dataset (K562 total RNA and mouse ES cell). These exons currently fail to be detected by the *CryptEx* algorithm.

No cryptic splicing events were shared between human and mouse as previously reported. Note that to report overlap with Ling and colleagues (datasets 1 and 2), the raw data was unsuitable for our cryptic exon discovery pipeline due to a lack of biological replicate samples. Instead the sequence data was aligned and the splice junctions generated by the aligner were used to classify previously reported cryptic exons.

In contrast, while a large number of novel splicing events were observed in the FUS depletion datasets, our algorithm only classified 3 in mouse and 1 in human as cryptic exons. FUS depletion was not observed to produce any cassette-like cryptic exons in either species. Figure S1 visualises the full unclassified output of the pipeline and illustrates the diversity of splicing alterations that occur upon RBP depletion. Figures S2 and S3 comprise of screenshots of every reported cryptic exon from the IGV browser (Robinson et al., 2011) in mouse and human respectively. Supplementary Tables 1 and 2 (ST1/ST2) list the coordinates of each cryptic exon along with the results of each experiment.

### 2.2 Cryptic exons are bound by TDP-43

For the remainder of our study, cryptic exons were grouped into unions of all cassette-like exons and extension events discovered in human and mouse, totalling 95 human and 52 murine cryptic exons. We then explored whether TDP-43 binding could explain the observed splicing changes in RNA-Seq data, as observed by Ling and colleagues. We took two complementary and genome-wide approaches: (i) searching for enriched motifs in the RNA sequence including and surrounding the cryptic exons and (ii) correlating the positions of cryptic exons with TDP-43 protein-RNA interaction data.

TDP-43 can repress or enhance the inclusion of a given exon by either binding within or adjacent to the exonic sequence. Hence, for our motif search, we flanked cryptic exon sequences by 100 nucleotides on either side. UG-rich motifs were found to be enriched in both mouse and human cryptic exons using two different algorithms: *MEME* (Fig 2A) and *HOMER* (S4). Of the 52 mouse cryptic exons 29 had a run of UG up to 40 nucleotides in length. Similarly, human cryptic exons were enriched in a UG motif but not in a continuous manner. By comparing the frequencies of 16 possible dinucleotides between the flanked cryptic exon sequence and the sequence of the adjacent intron either up or downstream of the cryptic-containing intron we were able to resolve the enrichment of UG dinucleotides. Figure 2D shows frequencies of the 16 different possible dinucleotides. UG and GU were enriched in flanked cryptic exon sequence in both human (fold change GU = 1.53; UG = 1.48; *P* < 10^−50^; proportion test) and mouse (fold change GU = 2.14; UG = 1.85; *P* < 10^−50^; proportion test).

**Figure 2.**
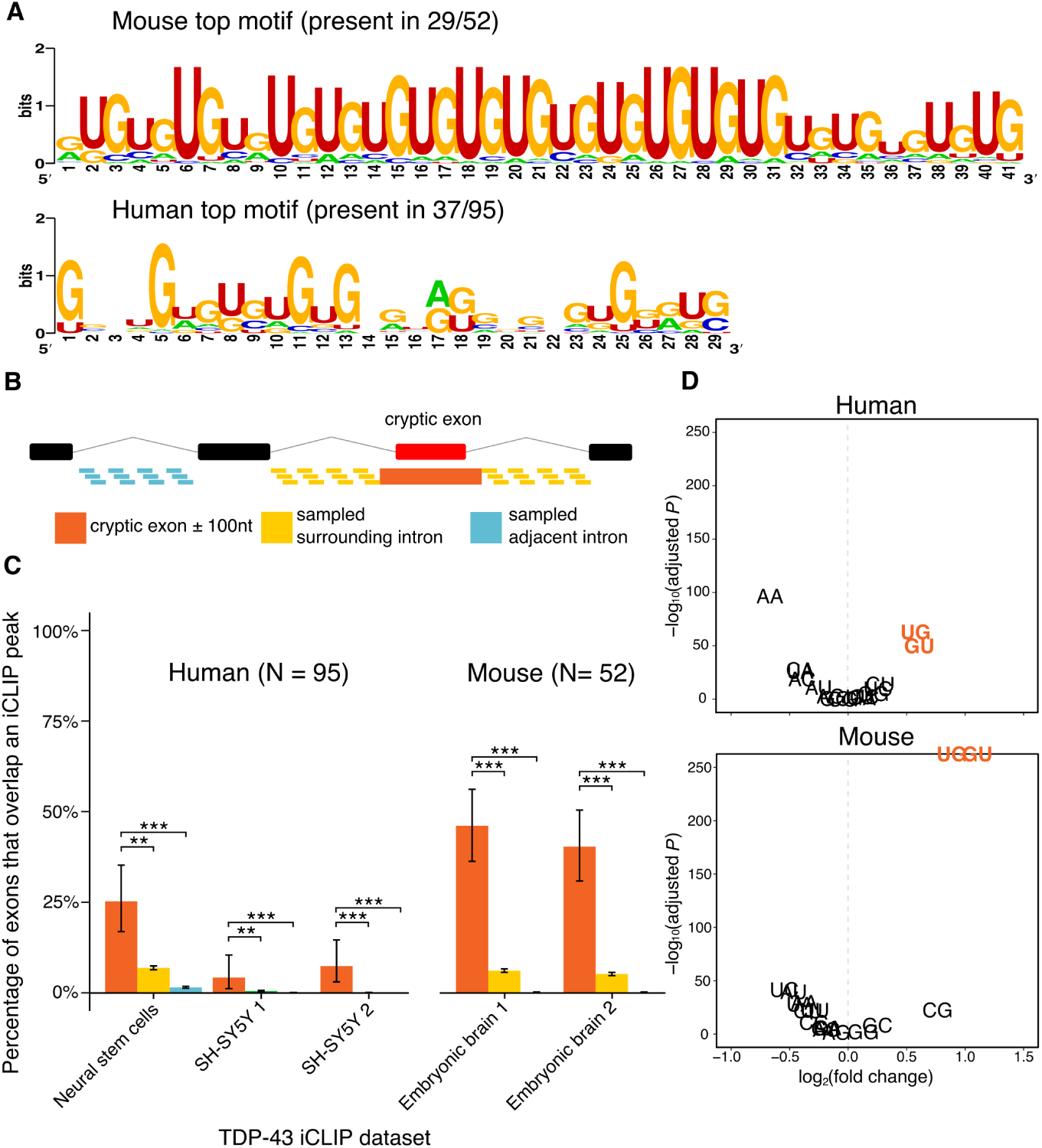
Evidence of TDP-43 binding cryptic exons.A– Results of *MEME* motif search. Only the motif with the greatest enrichment compared to background sequence is presented. B– Schematic of iCLIP peak enrichment test. For illustration a cassette-like cryptic exon (green box) is shown between two annotated exons (black boxes) separated by intronic sequence (black lines). The proportion of the group of cryptic exons flanked either side by 100 nucleotides (orange) that overlap at least one iCLIP peak is compared to the proportion of overlaps in a group of length matched sequences from either the surrounding intron (yellow) or an adjacent intron (blue) each randomly sampled 100 times per gene. C– iCLIP peak overlap enrichment for the 95 human cryptic exons found in either K562 cell TDP-43 depletion dataset and the 52 cryptic exons found in either mouse TDP-43 depletion dataset. D) Dinucleotide enrichment in the flanked cryptic exons compared to adjacent introns. Error bars denote 95% confidence intervals of the binomial distribution. ^*^*P* < 0.05, ^**^*P* < 0.001, ^***^*P* < 10^−16^. All P-values adjusted for multiple testing by Bonferroni method.

Individual nucleotide resolution UV crosslinking followed by immunoprecipation (iCLIP) allows for precise observation of which RNA species interact with a particular RNA-binding protein (Huppertz et al., 2014; Zarnack et al., 2013). We downloaded all publicly available TDP-43 iCLIP data from the iCount repository (http://icount.biolab.si/). We then intersected each set of iCLIP peaks with our cryptic exons, surrounded by 100 base pairs of flanking sequence either side. The proportion of overlapping cryptic exons was compared to the proportion of overlap between iCLIP peaks and two classes of null exon; the first created from the surrounding intronic sequence outside of each flanked cryptic and the second from an adjacent intron (see Figure 2B). If TDP-43 binding was uniform throughout an intron or throughout a gene then we would expect to see similar proportions of overlap in each. However, both species show an enrichment in TDP-43 binding peaks specific to the cryptic exons in every iCLIP dataset used, with as much as 25% of human cryptic exons and 50% of mouse cryptic exons overlapping at least one iCLIP peak each (both species: *P* < 10^−16^, proportion test).

### 2.3 Cryptic exon recognition is unrelated to the binding of transposable elements by TDP-43

Motivated by previous findings of cryptic exons associated with hnRNP C depletion and caused by exonisation of antisense Alu elements (Kelley et al., 2014; Zarnack et al., 2013), we investigated whether TDP-43 induced cryptic exons preferentially overlap specific families of transposable elements and/or class of repetitive sequences. Transposable/repeat elements annotations were obtained using the *RepeatMasker* software (Smit, AFA, Hubley, R & Green, P., 2015), and these features were split by family and orientation. Although Alu elements are a subfamily within the primate SINE element family, we included them separately given the prior hnRNP C result. Control regions were obtained by sampling similar length exons either within the same intron, or within adjacent introns. Figure 3 shows only modest enrichment patterns in the TDP-43 depletion datasets. This contrasts with hnRNP C depletion, which shows a striking enrichment of antisense SINE elements of which all are of the Alu type (*P* < 10^−16^, proportion test), a result consistent with previous analyses of dataset 9.

**Figure 3.**
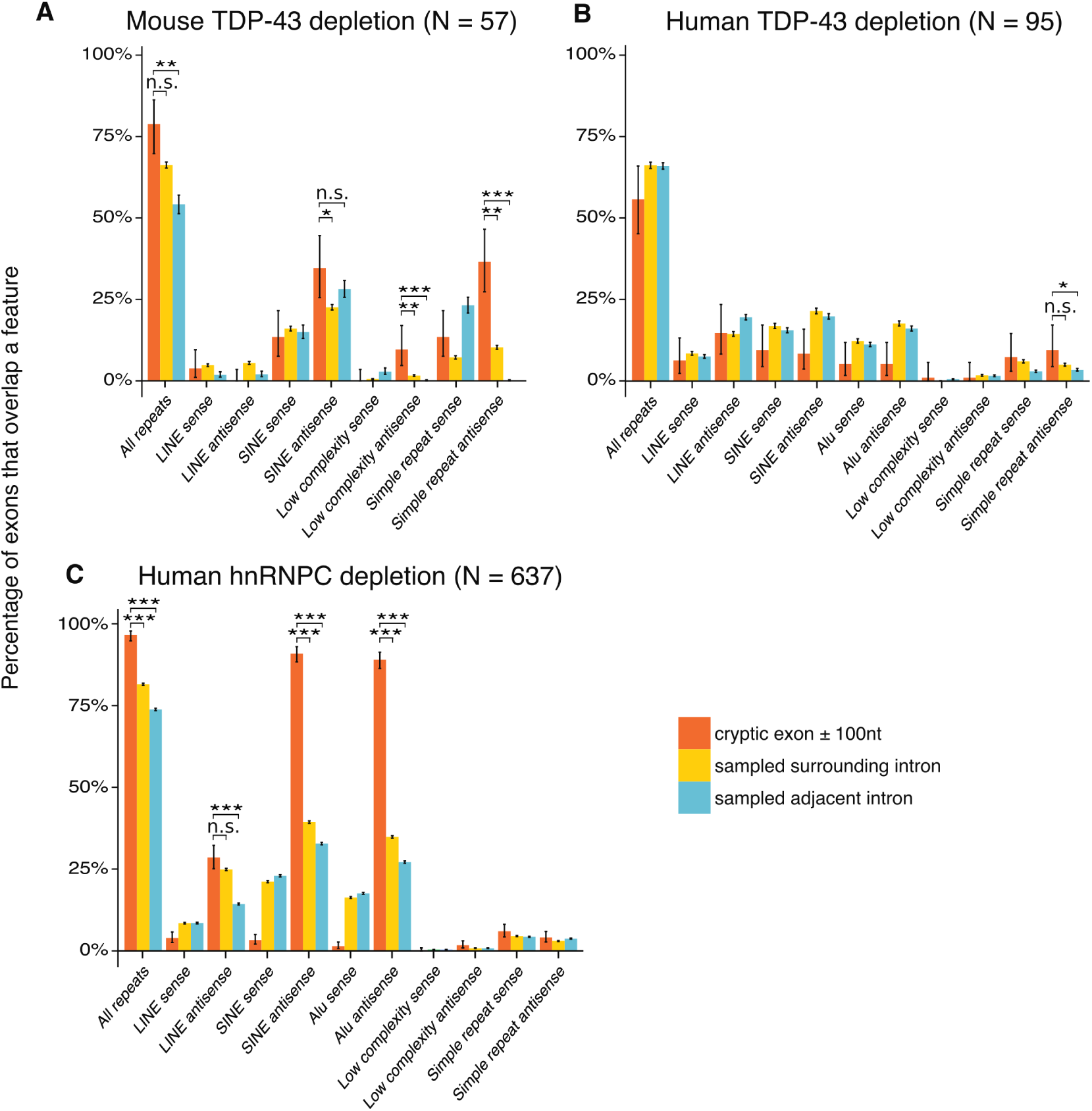
Cryptic exons in transposable elements. Overlap between different families of repetitive element with lists of exons, separated by orientation. A– Mouse TDP-43 depletion. B– Human TDP-43 depletion. C– Human hnRNP C depletion. The proportion of the cryptic exons that contain a particular element are shown in orange. Length matched random samples from the surrounding intron (yellow) and adjacent introns (blue) are used as controls. LINE: Long Interspersed Nuclear Element; SINE: Short Interspersed Nuclear Element. ^*^: *P* < 0.05, ^**^: *P* < 0.001, ^***^: *P* < 10^−16^. All P-values corrected for multiple testing with Bonferroni method.

### 2.4 Cryptic exons are poorly conserved and generate premature stop codons

We then quantified the extent of evolutionary conservation of cryptic exons using the multiple species alignment conservation scores generated by PhyloP (Pollard et al., 2010). We calculated mean PhyloP conservation scores per exon for the cryptic exons and compared them to both the annotated exons and the randomly sampled intronic sequences from the same genes. We found no difference between cryptic exons and matched intronic sequences (Figure 4A), and a much lower conservation level than adjacent annotated exons.

We also investigated the consequences of inclusion of cryptic exons on translation of the transcript. Potential outcomes for each gene are: (i) a functional transcript, (ii) a premature termination code (PTC) or (iii) a frameshift variant. We compared our estimates to random simulations where the identity of the included exon has been permuted 1000 times. Our results were consistent with the null, with 66% of cryptic exons leading to a frameshift due to length mismatches and less than 10% of cryptic exons predicted to create functional transcripts (Figure 4B).

**Figure 4.**
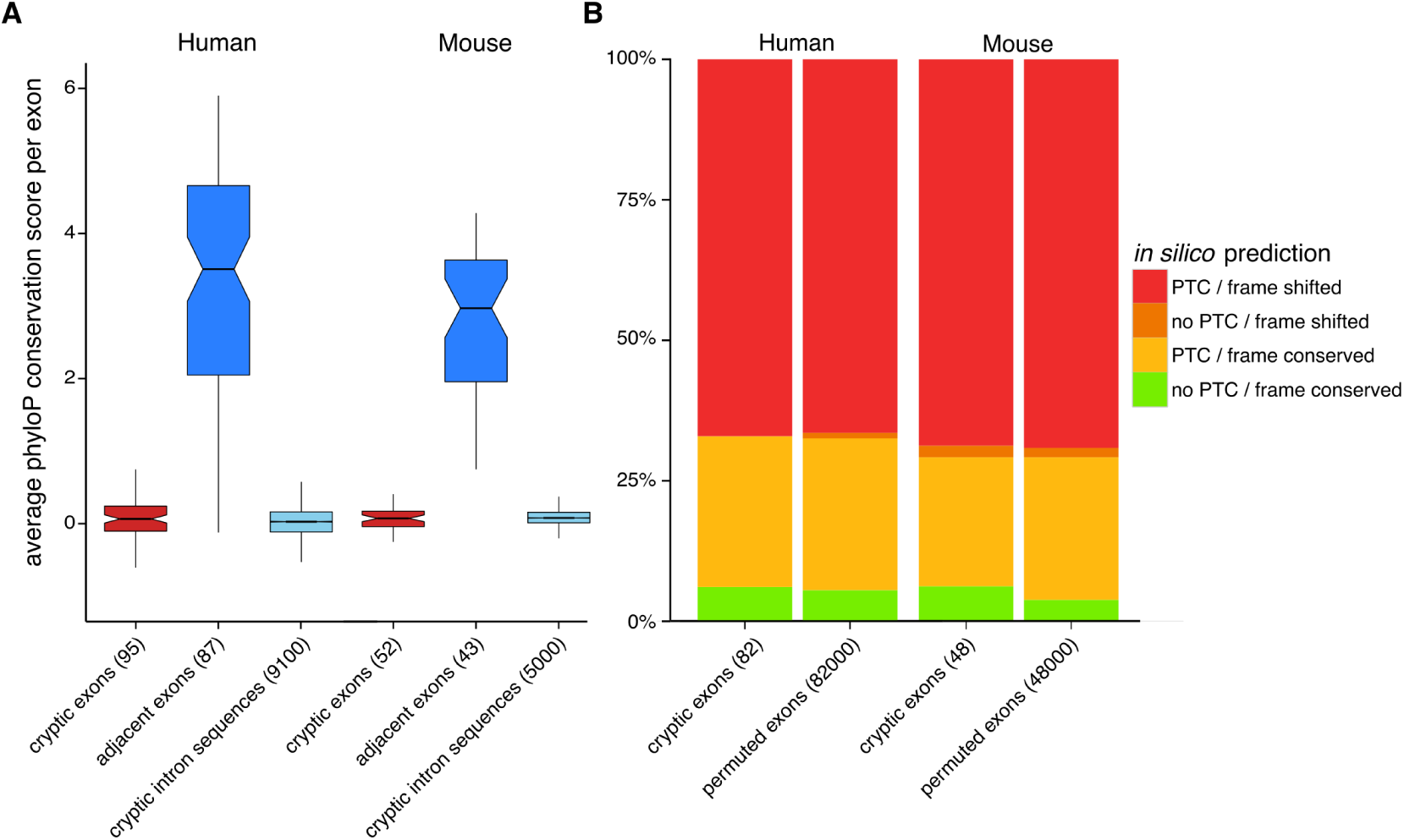
Conservation and premature termination codon analysis.A– Per exon average PhyloP conservation scores in cryptic exons, adjacent exons within the same set of genes (when available) and randomly sampled sequences from the cryptic containing intron. Box plots show first quartile, median and third quartile with the notches representing the 95% confidence interval of the median. Whiskers represent the minimum and maximum values that fall within 1.5 times the interquartile range. B- The functional impact of cryptic exon inclusion on the host transcripts in human and mouse. Colours indicate the category of prediction and box size indicates the proportion of the total group of exons in each category. Categories from top to bottom: at least one premature termination codon (PTC) introduced and frame shifted (red); no PTCs introduced but frame shifted (orange); PTCs introduced but frame conserved (yellow); no PTCs introduced and frame conserved (green). For each species there is a corresponding set of null exons where the central exon has been permuted 1000 times.

### 2.5 Cryptic exon containing genes are downregulated

We then investigated, in datasets 3-6, whether genes containing cryptic exons showed a specific pattern of altered expression. We calculated the proportion of the cryptic exon containing genes in each dataset that were differentially expressed at a FDR of 10%. We compared this with the proportion of differential expression of all genes with an expression level at or greater than the lowest expressed cryptic exon found in that dataset. Figure 5A shows the number of differentially expressed genes in each dataset as a proportion of the total, separated by direction. In all four TDP-43 depletion datasets, the cryptic exon containing genes as a group are more likely to be significantly downregulated compared to the genome-wide proportion (*P* < 0.001, hypergeometric test).

**Figure 5.**
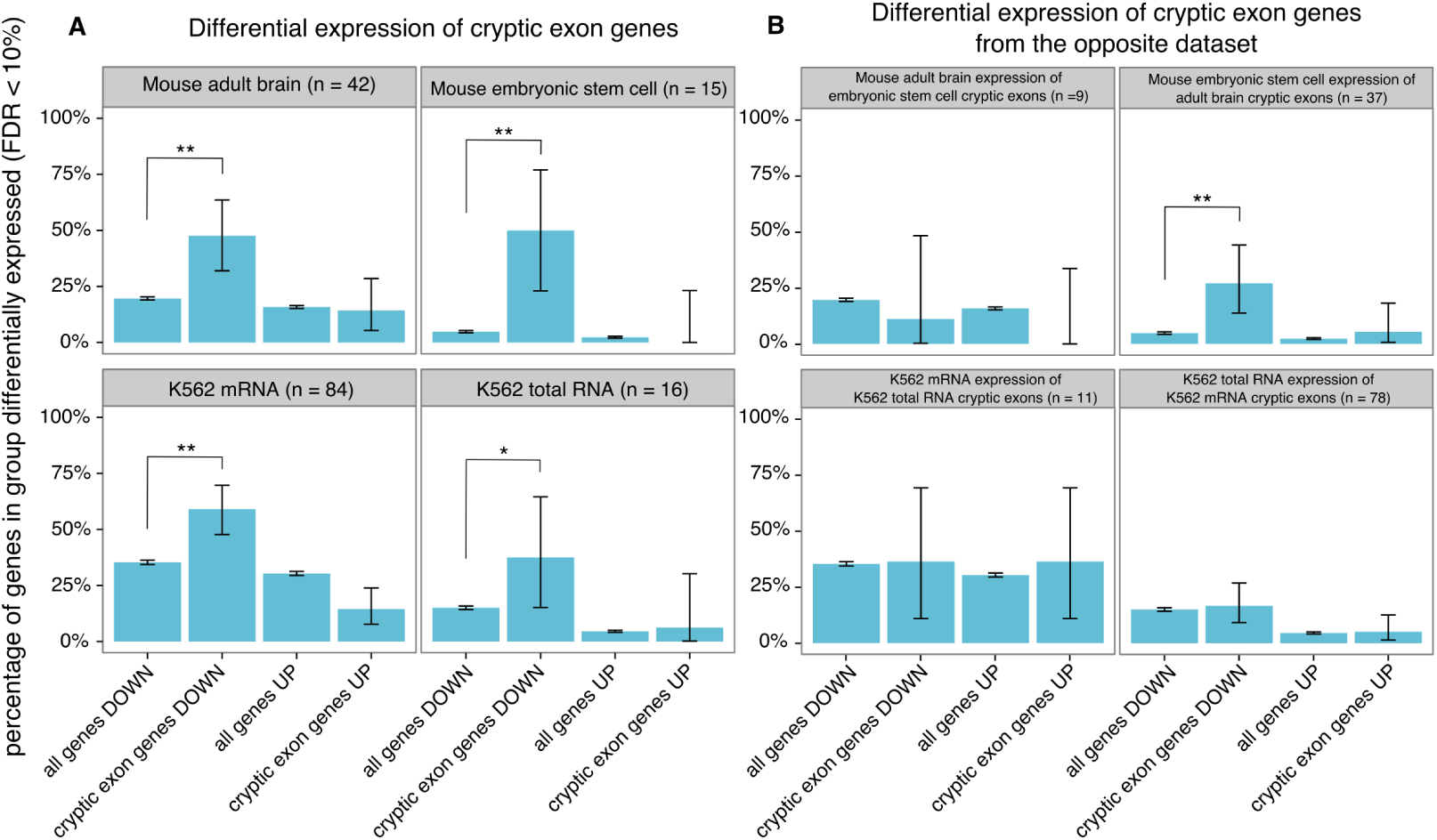
Differential expression of cryptic exon genes. A– Significantly differentially expressed genes between TDP-43 depletion and control samples (false discovery rate = 10%) in datasets 3-6. Comparisons were made between all genes tested with an expression level at or greater than the lowest expressed cryptic exon (“all genes”) and the genes where cryptic exons were discovered (“cryptic exon genes”) with each group divided by direction of change. B- As above but for cryptic exon genes from the other dataset of the same species. Error bars show 95% confidence intervals for each proportion. ^*^: *P* < 0.05, ^**^: *P* < 0.001. All P-values adjusted by Bonferroni correction.

Furthermore, we performed the same analysis for each dataset with the cryptic exon containing genes that were only found in the other dataset of the same species (Figure 5B). Surprisingly, in the mouse ES cell dataset 6 there was an enrichment of downregulated genes that contain cryptic exons only detectable in the mouse adult brain dataset 5 (*P* < 0.001, hypergeometric test). Visual inspection of these 10 introns in the mouse ES cell data (Figure S2) suggests that 7 of them may harbour cryptic exons in the ES cell data that are currently undetectable by the *CryptEx* algorithm.

### 2.6 Human cryptic exons are driven by the recognition of strong splice sites that are normally repressed

Whereas cassette-like cryptic exons appear as separate exons distinct from their surrounding exons, extension events must rely on a switch from a canonical splice site to a newly accessible splice site. We hypothesised that these extension events result from competition between two splice sites upon TDP-43 depletion. This would require the sequence of and around the cryptic splice site to be similarly recognisable to the spliceosome. Using the *MaxEnt* statistical model to score splice sites by comparing their DNA sequences with constitutive observed canonical sequences (Yeo et al., 2004), we scored the 5′ and 3′ splice sites of our cryptic exons and compared them with the scores of the surrounding canonical splice sites. The model compares splice sites from annotated exons with so-called decoy splice sites that retain the consensus AG/GT at the 3′ or 5′ splice site respectively. Therefore we also scored randomly generated sequences which retained the consensus AG/GT positions. Figure 6 shows the scores for both the 3′ and 5′ splice sites for each class of human cryptic exons. Although the canonical splice sites were on average stronger than their corresponding cryptic splice site (*P* < 0.05, paired t-test), the majority had scores far greater than those from random sequence, suggesting that they are able function as real, albeit weaker, splice sites when TDP-43 is depleted.

**Figure 6.**
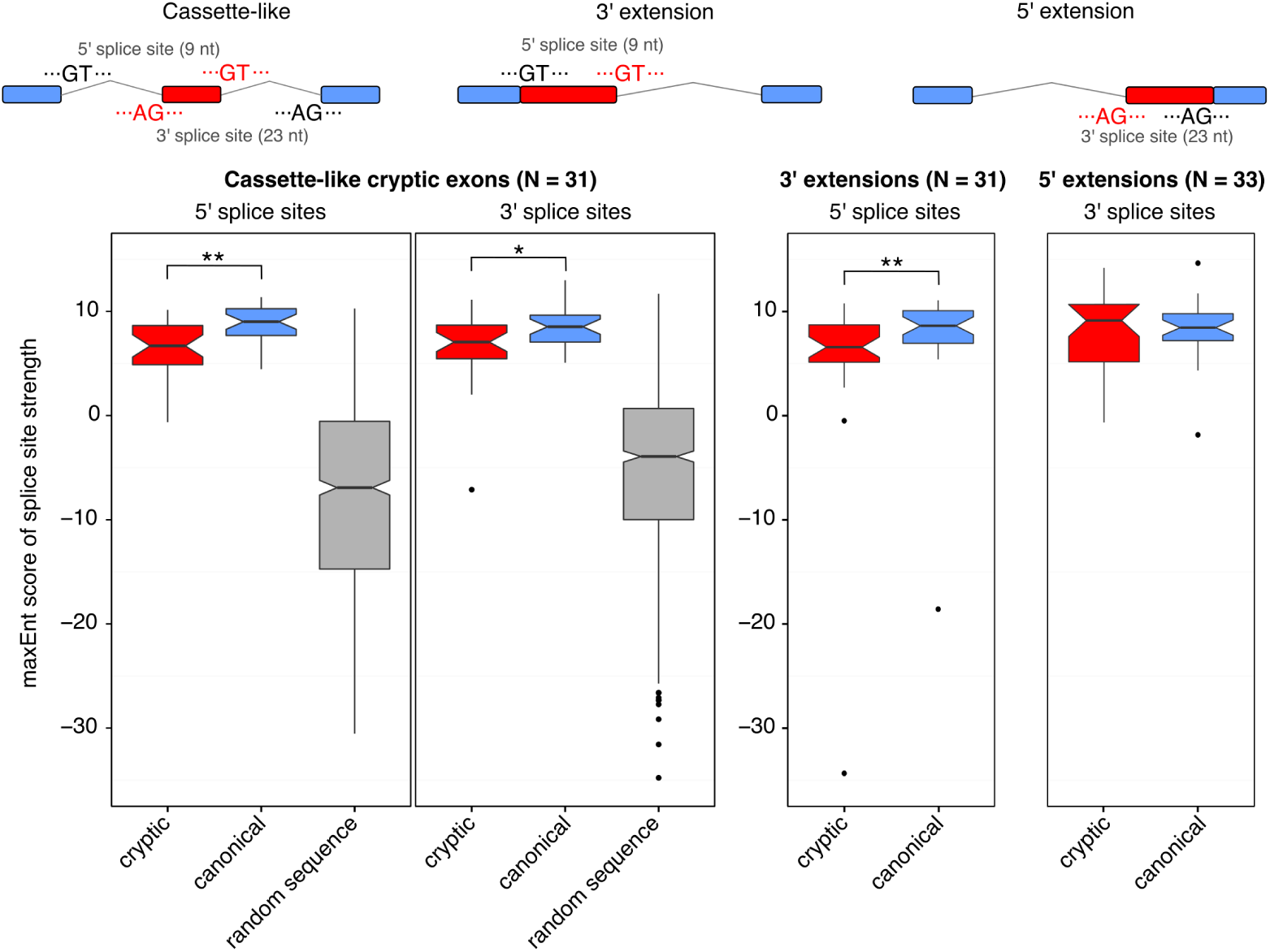
Scoring cryptic splice sites against canonical splice sites. Cassette-like cryptic exons have 5′ and 3′ splice sites that are recognised by the spliceosome under TDP-43 depletion. 5′ and 3′ splice site scores are plotted separately. Cryptic splice sites are shown in red, canonical splice sites are shown in blue. Random sequences with AG or GT consensus are plotted in grey. Cryptic extensions are the result of a cryptic splice site competing with the canonical splice site. 5′ extensions result from cryptic 3′ splice sites whereas 3′ extensions result from cryptic 5′ splice sites. Box plots show first quartile, median and third quartile with the notches representing the 95% confidence interval of the median. Whiskers represent the minimum and maximum values that fall within 1.5 times the interquartile range. Outliers are plotted as black dots. Cryptic splice sites are compared to canonical splice sites with paired t-tests.^*^: *P* < 0.05, ^**^: *P* < 0.001.

### 2.7 Cryptic exons are bound by other RNA binding proteins

Proteomic studies have demonstrated that TDP-43 interacts with a number of RNA-binding proteins (RBPs), including multiple members of the heterologously expressed ribonucleoprotein (hnRNP) family and other splicing factors (Blokhuis et al., 2016; Freibaum et al., 2010; Ling et al., 2010). The splicing of specific annotated exons has been shown to depend on the interaction of TDP-43 with multiple splicing factors (Mohagheghi et al., 2016). We hypothesised that some cryptic exons may be activated indirectly through TDP-43’s interaction with different RBPs. Van Nostrand and colleagues have performed eCLIP, a higher throughput modification of the iCLIP protocol, on 73 different RBPs including TDP-43 and FUS (Van Nostrand et al., 2016). The experiments were carried out in 2 human cell lines (K562 and HepG2) with 29 of the RBPs being tested in both cell lines. We performed the same overlap analysis between our human cryptic exons and each set of eCLIP peaks, using the same two sets of control sequences as before. Each eCLIP experiment was performed in duplicate. This gives each RBP four possible enrichment results using a proportion test. For each RBP, the highest P-value from the four tests was reported and corrected for multiple testing using a strict Bonferroni approach. Only proteins with a resulting *P* < 0.05 are reported. Unsurprisingly TDP-43 had the highest number of overlapping exons (*p* < 10^−22^; proportion test), followed by U2AF65, TIA1, SRSF7, U2AF35, PPIG, SRSF1 and IGF2BP1. Figure 6A shows the overlap between the different RBPs and the human cryptic exons. Figure 6B shows the overlap between each gene (columns) against each RBP (rows). Hierarchical clustering was performed on the RBPs. The three largest clusters consist of TDP-43 alone, the U2 snRNP binding proteins U2AF35 and U2AF65, and a third cluster containing the other proteins.

**Figure 7.**
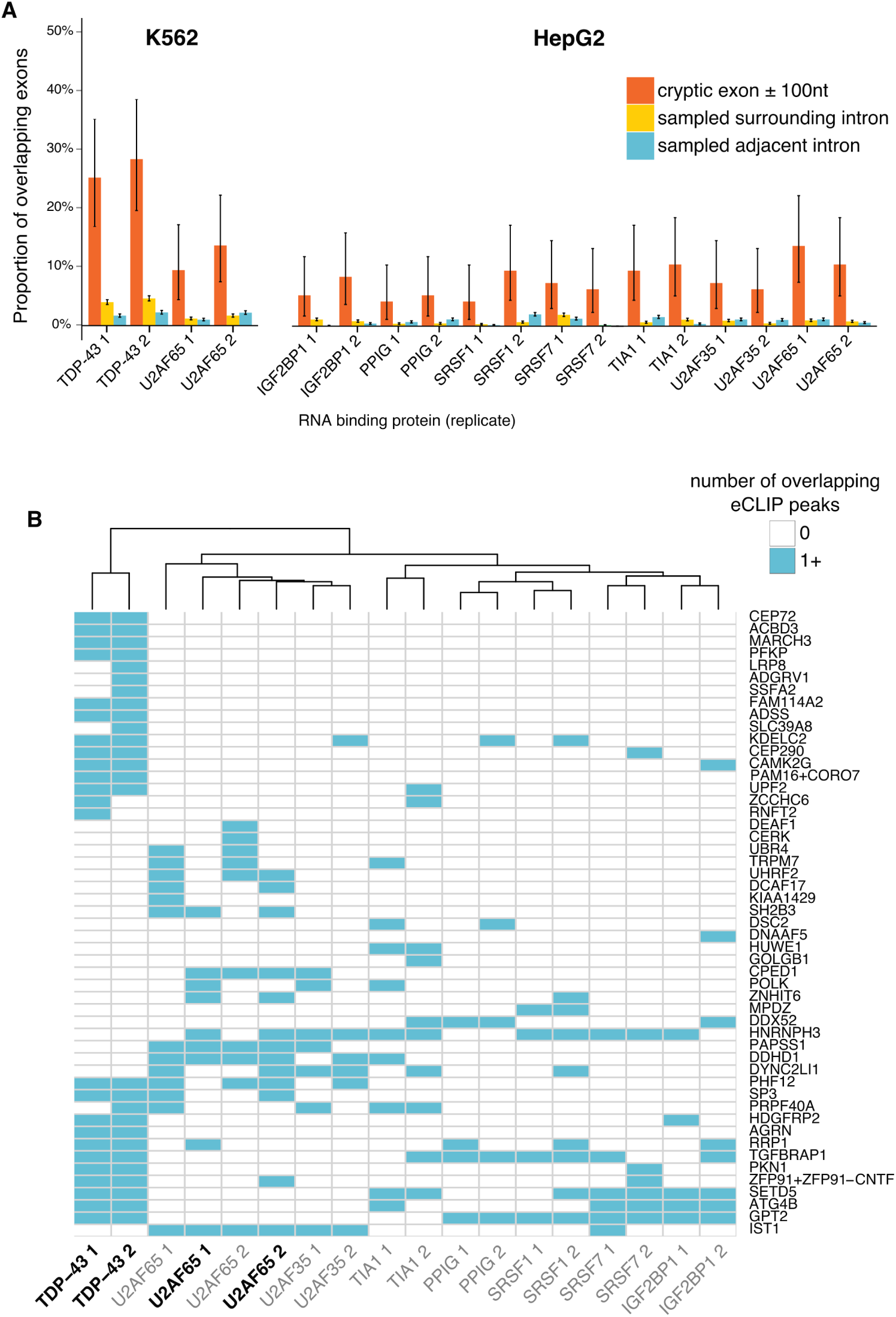
Mining of ENCODE eCLIP data in K562 and HepG2 cells. A– RNA-binding proteins (RBPs) with significant (*P* < 0.05) proportion of overlapping cryptic exons (red) compared to sampled intronic sequence from the same (green) and adjacent introns (blue). B- Comparison of eCLIP datasets from different RNA binding proteins (columns) showing the overlap with the cryptic exons (rows). RBPs in bold typeface are from K562 cells whereas those in regular typeface are from HepG2 cells.

## 3 Discussion

We have designed an analytical strategy to identify cryptic splicing that takes advantage of biological replicates in RNA sequencing data. We have applied this tool to a set of human and murine TDP-43 depletion datasets, as well as datasets that deplete hnRNP C or FUS. Our results are consistent with the previous findings that depletion of TDP-43 or hnRNP C leads to the inclusion of novel cryptic exons in both human and mouse. Although FUS undoubtedly plays an important role in splicing and mRNA stability and shares a number of targets with TDP-43 (Lagier-Tourenne et al., 2012), the low number of cryptic exons observed due to FUS depletion suggests that it does not play a major role in cryptic splicing and is a key point of differentiation with TDP-43.

Further examination of TDP-43 linked exons suggests they tend to possess the necessary UG-rich sequence elements to be bound by TDP-43 and using iCLIP data we observed that a subset of the cryptic exons are bound by TDP-43 *in vivo*. We went on to investigate the origins of these TDP-43 bound cryptic exons, as has been done for the targets of hnRNP C. We observed that unlike hnRNP C linked cryptic exons, which invariably originate from antisense Alu elements, TDP-43 linked cryptic exons do not originate from any single family of transposable element. Furthermore their sequences show very low species conservation, akin to random intronic sequence, but remarkably they contain splice sites very close in strength to those of their adjacent annotated exons. Our differential expression analysis suggests that the bulk of cryptic exon containing genes are significantly downregulated upon TDP-43 depletion. We hypothesize that this is due to nonsense mediated decay of the inclusion transcript, which we predict would occur in over 90% of cryptic exon containing transcripts. Consistent with this possibility, a recent proteomic study of TDP-43 depletion in human SH-SY5Y cells (Štalekar et al., 2015) that protein levels were changed for 3 of the 95 human cryptic exon containing genes. Two, *HUWE1* and *GOLGB1* had protein levels that were 8% and 31% of the control cells respectively whereas the third, *HNRNPH3* was found to be 7-fold increased under TDP-43 depletion. Interestingly, the cryptic exon discovered in hnRNP H3 falls upstream of the start codon whereas those found in *HUWE1* and *GOLGB1* are predicted to trigger NMD by inclusion into the coding sequence (see Supplementary Table 1). Another gene with cryptic splicing seen in both K562 datasets and in the initial Ling HeLa data, *AGRN*, was recently shown to be decreased at the protein level in the cerebrospinal fluid of ALS patients compared to healthy controls and other neurological diseases (Collins et al., 2015). Correct splicing of *AGRN* has shown to be crucial for the formation of the neuromuscular junction (Ruggiu et al., 2009).

Our current understanding of TDP-43’s role in cryptic splicing is that of a safeguard against the inclusion of potentially damaging intronic sequence into transcripts. However, the relationship between the UG-rich sequences and the strong 5′ and 3′ splice sites and their changes over evolutionary time are unknown as we observed no conserved cryptic exons between human and mouse.

Using publically available ENCODE eCLIP data, we identified a number of RNA binding proteins that also bind subsets of human cryptic exons under normal conditions, that is, in the presence of TDP-43. It is unsurprising that the splicing factors U2AF35 and U2AF65 are enriched as they preferably bind pyrimidine-rich 3’ splice site sequences which all cryptic exons appear to possess. That only 10-15% of cryptic exons show U2AF35/65 binding may be due to competition from TDP-43 in a manner similar to that seen between hnRNP C and U2AF65. IGF2BP1 has been reported as binding to TDP-43 in HEK293T and HeLa cell extracts (Freibaum et al., 2010; Ling et al., 2010), whereas SRSF7 was reported as binding to TDP-43 in mouse N2A cells (Blokhuis et al., 2016). None of the observed proteins have been reported to change their protein level in response to TDP-43 depletion (Štalekar et al., 2015).

As the majority of cryptic exons are predicted to lead to nonsense mediated decay of the inclusion transcript it seems peculiar that we can observe these transcripts at all. We hypothesise that cryptic splicing may be much more widespread than can be observed by RNA sequencing due to the highly efficient nature of nonsense mediated decay. Over half of all cryptic exon genes are significantly downregulated in each dataset (Fig 2B), suggesting that cryptic exon inclusion may be a key mechanism in the widespread changes in RNA expression that occur upon TDP-43 depletion.

Two genes, *ATG4B* and *GPSM2*, have previously been demonstrated to have cryptic exon inclusion RNA transcripts in ALS patient brain samples, suggesting a role for cryptic splicing in disease (Ling et al., 2015). Our analysis also identified a cryptic exon in *ATG4B* in human cells, but not *GPSM2*; however we did not analyse human brain data. By expanding the list of cryptic exons, it will be interesting to explore whether these are also dysregulated in ALS patient brains. However, such analysis may prove challenging owing to the likely small concentrations of RNA originating from diseased cells in brain homogenate and the likelihood of degradation by NMD. Alternate strategies may involve mass spectrometry screens for the subset of cryptic exon containing genes that escape the NMD process, for example because the cryptic exon is in frame. Such proteins may represent useful biomarkers for ALS.

In conclusion, we confirm the presence of cryptic exons after TDP-43 depletion and show they have a negative impact on the genes they reside in, leading to decreased expression levels. Further work is warranted to determine the relevance of cryptic exons to ALS and FTD pathogenesis.

## 4 Methods

### 4.1 Data preparation

Table 2 lists all the public data used in this study. For all RNA-seq data we first performed adapter and quality trimming (Phred score *>* 20) with *Trim Galore* (0.4.1) on the FASTQ files before aligning to either the human (hg38) or mouse (mm10) reference genome with *STAR* (2.4.2a). The resulting BAM file was sorted and PCR duplicate marked with *NovoSort* (1.03.09). Processed iCLIP peaks data was downloaded from the iCOUNT server (http://icount.biolab.si/). Both the human and mouse TDP-43 iCLIP data have been previously published (Rogelj et al., 2012; Tollervey et al., 2011).

**Table 2.**
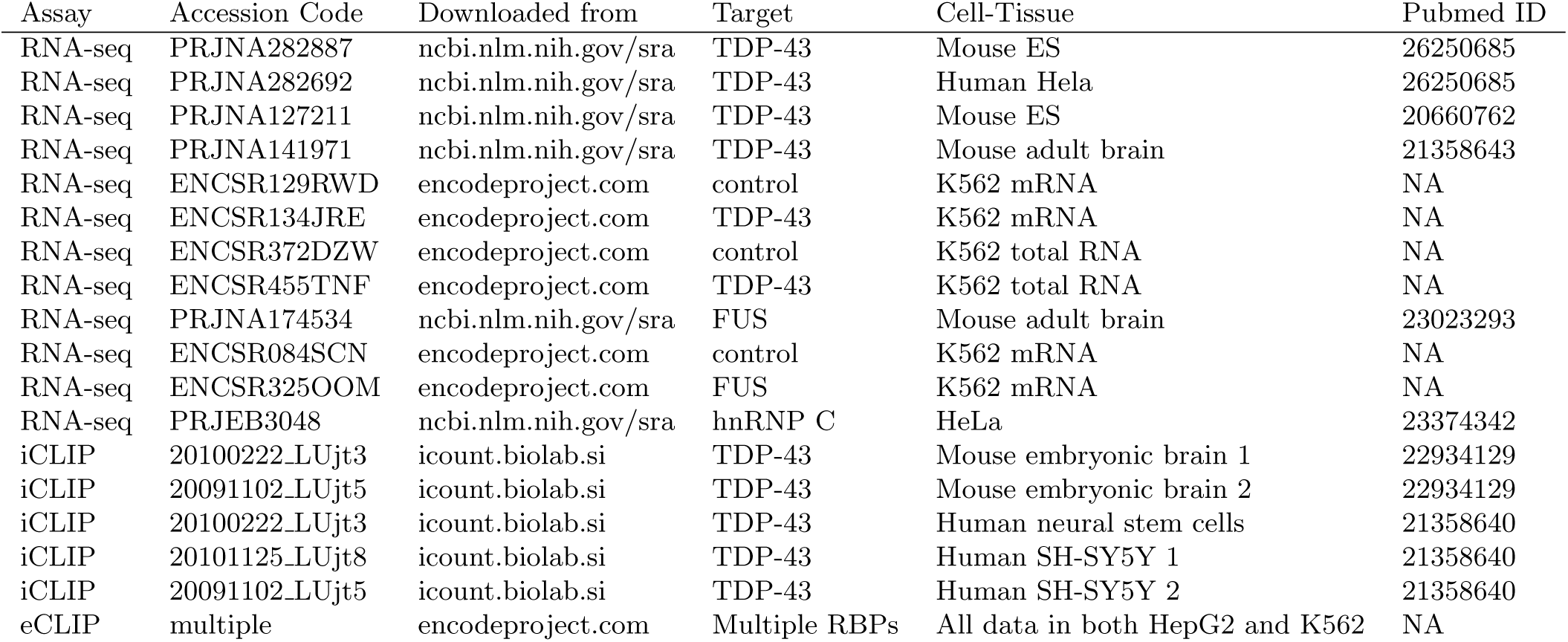
List of accessions. For the ENCODE RNA-seq libraries, the control and target depletion samples are listed under separate accessions.

Processed eCLIP data (previously described by (Van Nostrand et al., 2016)) was downloaded from the ENCODE project. The narrowPeaks bed format was used with the first nucleotide of the cluster defined as the peak. Peak coordinates from iCLIP and eCLIP were converted to the hg38 and mm10 builds using the *LiftOver* tool from UCSC.

### 4.2 Cryptic splicing definition

Splicing aware alignment software such as *STAR* (Dobin et al., 2013) cut short reads that originate from a spliced transcript and align the pieces separately, marking the distance between them as a splice junction. Splice junctions can be used to reaffirm known splicing patterns or infer novel splicing. We define cryptic splicing as the emergence or relative increase in splice junctions that splice from known splice sites to unannotated positions within introns. This increase correlates with the depletion of a particular RNA binding protein. Different repositories have different levels of proof for annotating exons but we define an annotated exon as one listed in the Ensembl list of transcripts (release 82).

### 4.3 Cryptic splicing discovery with the *CryptEx* pipeline

Due to the diversity in the quality of published RNA-seq data, the cryptic splicing discovery pipeline was designed to be used on any RNA-seq library, whether single or paired end, stranded or unstranded, total RNA or polyA-selected RNA. This flexibility results in a large number of false positive hits which have to be aggressively filtered downstream. The initial cryptic exon discovery pipeline was written in Bash, using *SAMTools* (Li et al., 2009) (version 1.2) and *BEDTools* (Quinlan and Hall, 2010) (version 2.25.0). All code for downstream processing and filtering of cryptic hits was written in the R language (version 3.1.1) using the *Biostrings, data.table, DEXSeq, dplyr, GenomicRanges, ggplot2, gridExtra, optparse, plyr, stringr,* and *tidyr* packages. All the code for reproducing this paper is available in a GitHub repository (https://github.com/jackhump/CryptEx).

In order to discover all possible splice junctions that travel into the intron we first extracted all spliced reads from each aligned bam file using *SAMTools*, filtering out any secondary alignments. To extract only the spliced reads that overlap an annotated exon we then performed an intersection in *BEDTools* with a flattened list of exons, created using the dexseq prepare annotation.py Python script included with the *DEXSeq* package (Anders et al., 2012). An inverse intersection was then performed with the same exon list to retain only the spliced reads which do not bridge two annotated exons. The intronic mapping sections of each read were split off from the rest and retained. In each dataset all intronic mapping spliced reads from each sample were grouped together irrespective of condition. Split reads that were within 500bp of each other were merged into larger intervals, hereby referred to as tags. This ideally captures both the upstream and downstream splice junction to a central cryptic cassette exon. To keep only the tags that are splicing within the gene body another intersection was performed with a list of introns. This was generated from the same flattened exon file by an R script written by Devon Ryan (Ryan, 2014). The tags were then incorporated into the flattened list of exons. The reads that overlap annotated exons and tags were counted using HTSeq on the default settings, ignoring PCR duplicate reads (Anders et al., 2015). The read counts were used to calculate differential usage of each exon with DEXSeq.

All the cryptic tags with an adjusted P-value (false discovery rate) < 5% and a log_2_ fold change > |0.6| were extracted from the DEXSeq results table (see Figure S1). The splice junctions from the alignment of each sample were used to work out the coordinates of the canonical junction that spans the intron within which the cryptic tag is or isn’t spliced in control samples. Using splice junctions from the depletion condition samples, the upstream and downstream junctions that connect the adjacent annotated exons to the cryptic tag were re-discovered and quantified. Any cryptic example that did not have at least one upstream or downstream junction per sample or had fewer than ten canonical splice junctions was removed. These junctions were used to calculate per-condition mean Percent Spliced In (PSI) values which are a ratio of cryptic splicing over the sum of cryptic and canonical splicing (Katz et al., 2010). As a number of cryptic splicing events are present at a low level in control samples, delta PSI values were created for both upstream and downstream splicing for each tag. This is the difference in PSI between the depletion samples and the control samples. Any cryptic tag that had either an upstream or downstream delta PSI < 5% was removed.

### 4.4 iCLIP/eCLIP enrichment

The coordinates of each cryptic tag were flanked by 100 base pairs on either side to capture binding around the putative splice sites. In order to compare the overlap between cryptic exons and RNA-protein binding peaks, two sets of null exons were created for comparison, which maintain the same length as their corresponding cryptic exon but sample either the intronic sequence outside of the flanked exon or that of the adjacent introns within the same gene if available. Overlaps between exons and iCLIP and eCLIP peaks were calculated using BedTools.

### 4.5 Motif enrichment analysis

FASTA sequence was generated for the cryptic exons flanked by 100 nucleotides either side and submitted to the *MEME* web tool under the default settings (Bailey et al., 2009). The analysis was repeated using the *HOMER* algorithm on RNA mode (Heinz et al., 2010). Motifs were created using *WebLogo* (Crooks et al., 2004). Frequencies of the 16 possible dinucleotides were compared between flanked cryptic exon sequences with adjacent intron sequences from the same gene.

### 4.6 Transposable element enrichment

Lists of transposable elements in human and mouse (hg38 and mm10 respectively) were previously generated by the *RepeatMasker* tool and were downloaded from UCSC. Overlap between different transposable elements and the cryptic exons was calculated in each orientation using *BedTools*.

### 4.7 Conservation analysis

PhyloP compares the sequence alignments of multiple species to produce per base conservation scores (Pollard et al., 2010). Average conservation score per cryptic tag was calculated using *bigWigSummary* (UCSC) for both human and mouse data. The lists of splice junctions created by *STAR* when aligning each sample were used to identify the coordinates of the exons adjacent to the cryptic exon. The randomly sampled intronic sequence from the cryptic-containing intron was used as a negative control.

### 4.8 Protein prediction analysis

Any cryptic exon which did not fall within the coding sequence of a transcript was omitted. Splice junctions were used to determine the upstream and downstream exons adjacent to each cryptic exon. These exons were matched to their corresponding annotated exon in the Ensembl transcript file for each species to work out the correct frame of translation. Nucleotide sequences for transcripts either including or excluding the cryptic tag were created. If the cryptic exon had been previously flagged as an extension then the entire continuous intronic sequence was inserted up to the remaining cryptic splice site. These transcripts were then translated in silico and defined as premature termination codon (PTC)-containing if the inclusion transcript contained a stop codon and as a frameshift if the sequence of the downstream exon no longer matched. A null distribution of PTC-containing or frame shifted transcripts was created by shuffling the identity of the central exon 1000 times.

### 4.9 Splice junction scoring

The strength of 5′ and 3′ splice sites was calculated for the human cryptic exons using *maxEnt* (Yeo et al., 2004). Higher scores indicate the increased log odds of a given splice site being a true splice site. The 5′ splice site is defined as the last 3 nucleotides of the upstream exon flanked by 6 intronic nucleotides, of which the first two are invariably GU. The 3′ splice site is defined as the last 20 intronic nucleotides of which the final two are invariably AG, flanked by the first 3 nucleotides of the downstream exon. The splice sites of annotated exons were used as a positive control. Randomly generated sequence with invariant AG or GT was used as a negative control. Paired t-tests were carried out to test the direction of change between the cryptic and annotated splice sites for each class of cryptic exon.

## 5 Acknowledgments

The authors would like to thank all members of the Plagnol, Fratta and Isaacs labs for their assistance and discussion. We are extremely grateful for the RNA-seq, iCLIP and eCLIP data made freely available by the Cleveland, Gravely, Wong, Ule and Yeo labs. J.H. is supported by a Medical Research Council and Brain Research Trust PhD studentship. The work of W.E. is supported by the Wellcome Trust (103760/Z/14/Z) and the Francis Crick Institute, which receives its core funding from Cancer Research UK (FC001002), the UK Medical Research Council (FC001002), and the Wellcome Trust (FC001002). P.F. is funded by a Medical Research Council/Motor Neuron Disease Association LEW Fellowship and by the NIHR UCLH BRC. A.M.I. is funded by the Medical Research Council, the Motor Neuron Disease Association and the European Research Council.

## 6 Contributions

J.H. and V.P. wrote all the code used to analyse the data with input from W.E.. J.H., V.P., P.F., and A.M.I. designed the experiments. J.H. wrote the manuscript with discussion and approval from all authors.

## References

Anders S, Pyl PT, and Huber W. 2015. HTSeq–a python framework to work with high-throughput sequencing data. Bioinformatics 31: 166–169.

Anders S, Reyes A, and Huber W. 2012. Detecting differential usage of exons from RNA-seq data. Genome Res. 22: 2008–2017.

Arnold ES, S-C L, Huelga SC, Lagier-Tourenne C, Polymenidou M, Ditsworth D, Kordasiewicz HB, McAlonis-Downes M, Platoshyn O, Parone PA, et al. 2013. ALS-linked TDP-43 mutations produce aberrant RNA splicing and adult-onset motor neuron disease without aggregation or loss of nuclear TDP-43. Proceedings of the National Academy of Sciences 110: E736–E745.

Bai Y, Ji S, and Wang Y. 2015. IRcall and IRclassifier: two methods for flexible detection of intron retention events from RNA-Seq data. BMC Genomics 16 Suppl 2: S9.

Bailey TL, Boden M, Buske FA, Frith M, Grant CE, Clementi L, Ren J, Li WW, and Noble WS. 2009. MEME SUITE: tools for motif discovery and searching. Nucleic Acids Res. 37: W202–8.

Blokhuis AM, Koppers M, Groen EJN, van den Heuvel DMA, Dini Modigliani S, Anink JJ, Fumoto K, van Diggelen F, Snelting A, Sodaar P, et al. 2016. Comparative interactomics analysis of different ALS-associated proteins identifies converging molecular pathways. Acta Neuropathol. 132: 175–196.

Bose JK, Wang IF, Hung L, Tarn WY, and Shen CKJ. 2008. TDP-43 overexpression enhances exon 7 inclusion during the survival of motor neuron pre-mRNA splicing. J. Biol. Chem. 283: 28852–28859.

Buratti E and Baralle FE. 2001. Characterization and functional implications of the RNA binding properties of nuclear factor TDP-43, a novel splicing regulator of CFTR exon 9. J. Biol. Chem. 276: 36337–36343.

Buratti E, Chivers M, Kralovicova J, Romano M, Baralle M, Krainer AR, and Vorechovsky I. 2007. Aberrant 5’ splice sites in human disease genes: mutation pattern, nucleotide structure and comparison of computational tools that predict their utilization. Nucleic Acids Res. 35: 4250–4263.

Chiang PM, Ling J, Jeong YH, Price DL, Aja SM, and Wong PC. 2010. Deletion of TDP-43 down-regulates tbc1d1, a gene linked to obesity, and alters body fat metabolism. Proc. Natl. Acad. Sci. U. S. A. 107: 16320–16324.

Collins MA, An J, Hood BL, Conrads TP, and Bowser RP. 2015. Label-Free LC-MS/MS proteomic analysis of cerebrospinal fluid identifies Protein/Pathway alterations and candidate biomarkers for amyotrophic lateral sclerosis. J. Proteome Res. 14: 4486–4501.

Crooks GE, Hon G, Chandonia JM, and Brenner SE. 2004. WebLogo: a sequence logo generator. Genome Res. 14: 1188–1190.

Deininger P and Prescott D. 2011. Alu elements: know the SINEs. Genome Biol. 12: 236.

Dobin A, Davis CA, Schlesinger F, Drenkow J, Zaleski C, Jha S, Batut P, Chaisson M, and Gingeras TR. 2013. STAR: ultrafast universal RNA-seq aligner. Bioinformatics 29: 15–21.

Eng L, Coutinho G, Nahas S, Yeo G, Tanouye R, Babaei M, Dörk T, Burge C, and Gatti RA. 2004. Nonclassical splicing mutations in the coding and noncoding regions of the ATM gene: maximum entropy estimates of splice junction strengths. Hum. Mutat. 23: 67–76.

Freibaum BD, Chitta RK, High AA, and Taylor JP. 2010. Global analysis of TDP-43 interacting proteins reveals strong association with RNA splicing and translation machinery. J. Proteome Res. 9: 1104–1120.

Heinz S, Sven H, Christopher B, Nathanael S, Eric B, Lin YC, Peter L, Cheng JX, Cornelis M, Harinder S, et al. 2010. Simple combinations of Lineage-Determining transcription factors prime cis-regulatory elements required for macrophage and B cell identities. Mol. Cell 38: 576–589.

Huppertz I, Ina H, Jan A, Andrea D, Easton LE, Sibley CR, Yoichiro S, Mojca T, Julian K, and Jernej U. 2014. iCLIP: Protein–RNA interactions at nucleotide resolution. Methods 65: 274–287.

Katz Y, Wang ET, Airoldi EM, and Burge CB. 2010. Analysis and design of RNA sequencing experiments for identifying isoform regulation. Nat. Methods 7: 1009–1015.

Kelley DR, Hendrickson DG, Tenen D, and Rinn JL. 2014. Transposable elements modulate human RNA abundance and splicing via specific RNA-protein interactions. Genome Biol. 15: 537.

Kim HJ, Kim NC, Wang YD, Scarborough EA, Moore J, Diaz Z, MacLea KS, Freibaum B, Li S, Molliex A, et al. 2013. Mutations in prion-like domains in hnRNPA2B1 and hnRNPA1 cause multisystem proteinopathy and ALS. Nature 495: 467–473.

Lagier-Tourenne C, Polymenidou M, Hutt KR, Vu AQ, Baughn M, Huelga SC, Clutario KM, Ling SC, Liang TY, Mazur C, et al. 2012. Divergent roles of ALS-linked proteins FUS/TLS and TDP-43 intersect in processing long pre-mRNAs. Nat. Neurosci. 15: 1488–1497.

Li H, Handsaker B, Wysoker A, Fennell T, Ruan J, Homer N, Marth G, Abecasis G, Durbin R, and 1000 Genome Project Data Processing Subgroup. 2009. The sequence Alignment/Map format and SAMtools. Bioinformatics 25: 2078–2079.

Li Y, Rao X, Mattox WW, Amos CI, and Liu B. 2015. RNA-Seq analysis of differential splice junction usage and intron retentions by DEXSeq. PLoS One 10: e0136653.

Ling JP, Pletnikova O, Troncoso JC, and Wong PC. 2015. TDP-43 repression of nonconserved cryptic exons is compromised in ALS-FTD. Science 349: 650–655.

Ling SC, Albuquerque CP, Han JS, Lagier-Tourenne C, Tokunaga S, Zhou H, and Cleveland DW. 2010. ALS-associated mutations in TDP-43 increase its stability and promote TDP-43 complexes with FUS/TLS. Proc. Natl. Acad. Sci. U. S. A. 107: 13318–13323.

Matera AG, Gregory Matera A, and Zefeng W. 2014. A day in the life of the spliceosome. Nat. Rev. Mol. Cell Biol. 15: 108–121.

McGlincy NJ and Smith CWJ. 2008. Alternative splicing resulting in nonsense-mediated mRNA decay: what is the meaning of nonsense? Trends Biochem. Sci. 33: 385–393.

Meili D, David M, Jana K, Julian Z, Luisa B, Laura F, Nenad B, Beat T, and Igor V. 2009. Disease-causing mutations improving the branch site and polypyrimidine tract: Pseudoexon activation of LINE-2 and antisense alu lacking the poly(t)-tail. Hum. Mutat. 30: 823–831.

Mercado PA, Ayala YM, Romano M, Buratti E, and Baralle FE. 2005. Depletion of TDP 43 overrides the need for exonic and intronic splicing enhancers in the human apoA-II gene. Nucleic Acids Res. 33: 6000–6010.

Mohagheghi F, Prudencio M, Stuani C, Cook C, Jansen-West K, Dickson DW, Petrucelli L, and Buratti E. 2016. TDP-43 functions within a network of hnRNP proteins to inhibit the production of a truncated human SORT1 receptor. Hum. Mol. Genet. 25: 534–545.

Neumann M, Sampathu DM, Kwong LK, Truax AC, Micsenyi MC, Chou TT, Bruce J, Schuck T, Grossman M, Clark CM, et al. 2006. Ubiquitinated TDP-43 in frontotemporal lobar degeneration and amyotrophic lateral sclerosis. Science 314: 130–133.

Pimentel H, Conboy JG, and Pachter L. 2015. Keep me around: Intron retention detection and analysis. ArXiv.

Pollard KS, Hubisz MJ, Rosenbloom KR, and Siepel A. 2010. Detection of nonneutral substitution rates on mammalian phylogenies. Genome Res. 20: 110–121.

Polymenidou M, Lagier-Tourenne C, Hutt KR, Huelga SC, Moran J, Liang TY, Ling SC, Sun E, Wancewicz E, Mazur C, et al. 2011. Long pre-mRNA depletion and RNA missplicing contribute to neuronal vulnerability from loss of TDP-43. Nat. Neurosci. 14: 459–468.

Quinlan AR and Hall IM. 2010. BEDTools: a flexible suite of utilities for comparing genomic features. Bioinformatics 26: 841–842.

Robinson JT, Helga T, Wendy W, Mitchell G, Lander ES, Gad G, and Mesirov JP. 2011. Integrative genomics viewer. Nat. Biotechnol. 29: 24–26.

Rogelj B, Easton LE, Bogu GK, Stanton LW, Rot G, Curk T, Zupan B, Sugimoto Y, Modic M, Haberman N, et al. 2012. Widespread binding of FUS along nascent RNA regulates alternative splicing in the brain. Sci. Rep. 2: 603.

Ruggiu M, Herbst R, Kim N, Jevsek M, Fak JJ, Mann MA, Fischbach G, Burden SJ, and Darnell RB. 2009. Rescuing Z agrin splicing in nova null mice restores synapse formation and unmasks a physiologic defect in motor neuron firing. Proceedings of the National Academy of Sciences 106: 3513–3518.

Ryan D. 2014. DEXSeq for intron retention. http://seqanswers.com/forums/archive/index.php/t-42420.html. Accessed: 2015-10-1.

Shan X, P-M C, Price DL, and Wong PC. 2010. Altered distributions of gemini of coiled bodies and mitochondria in motor neurons of TDP-43 transgenic mice. Proceedings of the National Academy of Sciences 107: 16325–16330.

Shiga A, Ishihara T, Miyashita A, Kuwabara M, Kato T, Watanabe N, Yamahira A, Kondo C, Yokoseki A, Takahashi M, et al. 2012. Alteration of POLDIP3 splicing associated with loss of function of TDP-43 in tissues affected with ALS. PLoS One 7: e43120.

Smit, AFA, Hubley, R & Green, P. 2015. RepeatMasker open-4.0. http://www.repeatmasker.org. Accessed: 2016-2-1.

Sorek R, Ast G, and Graur D. 2002. Alu-containing exons are alternatively spliced. Genome Res. 12: 1060–1067.

Sreedharan J, Blair IP, Tripathi VB, Hu X, Vance C, Rogelj B, Ackerley S, Durnall JC, Williams KL, Buratti E, et al. 2008. TDP-43 mutations in familial and sporadic amyotrophic lateral sclerosis. Science 319: 1668–1672.

Tollervey JR, Tomaž C, Boris R, Michael B, Matteo C, Melis K, Julian K, Tibor H, Nishimura AL, Vera v, et al. 2011. Characterizing the RNA targets and position-dependent splicing regulation by TDP-43. Nat. Neurosci. 14: 452–458.

Van Nostrand EL, Pratt GA, Shishkin AA, Gelboin-Burkhart C, Fang MY, Sundararaman B, Blue SM, Nguyen TB, Surka C, Elkins K, et al. 2016. Robust transcriptome-wide discovery of RNA-binding protein binding sites with enhanced CLIP (eCLIP). Nat. Methods 13: 508–514.

Vance C, Rogelj B, Hortobaágyi T, De Vos KJ, Nishimura AL, Sreedharan J, Hu X, Smith B, Ruddy D, Wright P, et al. 2009. Mutations in FUS, an RNA processing protein, cause familial amyotrophic lateral sclerosis type 6. Science 323: 1208–1211.

Vorechovsky I. 2006. Aberrant 3’ splice sites in human disease genes: mutation pattern, nucleotide structure and comparison of computational tools that predict their utilization. Nucleic Acids Res. 34: 4630–4641.

Vorechovsky I. 2010. Transposable elements in disease-associated cryptic exons. Hum. Genet. 127: 135–154.

Štalekar M, Yin X, Rebolj K, Darovic S, Troakes C, Mayr M, Shaw CE, and Rogelj B. 2015. Proteomic analyses reveal that loss of TDP-43 affects RNA processing and intracellular transport. Neuroscience 293: 157–170.

Yeo G, Gene Y, and Burge CB. 2004. Maximum entropy modeling of short sequence motifs with applications to RNA splicing signals. J. Comput. Biol. 11: 377–394.

Zarnack K, König J, Tajnik M, Martincorena In, Eustermann S, Stévant I, Reyes A, Anders S, Luscombe NM, and Ule J. 2013. Direct competition between hnRNP C and U2AF65 protects the transcriptome from the exonization of alu elements. Cell 152: 453–466.

